# Prevalence of *Mycoplasma genitalium* in different population groups: systematic review and meta-analysis

**DOI:** 10.1101/180422

**Authors:** Lukas Baumann, Manuel Cina, Dianne Egli-Gany, Myrofora Goutaki, Florian Halbeisen, Gian-Reto Lohrer, Hammad Ali, Pippa Scott, Nicola Low

**Affiliations:** Institute of Social and Preventive Medicine, University of Bern, Switzerland; Kirby Institute, University of New South Wales, Australia; University of Otago, New Zealand

## Abstract

**Background:** *Mycoplasma genitalium* is a common cause of non-gonococcal non-chlamydial urethritis and cervicitis. Testing of asymptomatic populations has been proposed, but prevalence rates in asymptomatic populations are not well established. We aimed to estimate the prevalence of *M. genitalium* in adults in the general population, in clinic-based samples, pregnant women, men who have sex with men (MSM) and female sex workers (FSW).

**Methods:** We searched Embase, Medline, IndMED, AIM and LILACS from 1 January 1991 to 12 July 2016 without language restrictions. We included studies with 500 participants or more. We screened and selected studies and extracted data in duplicate. We examined eligible studies in forest plots and conducted random effects meta-analysis to estimate prevalence, if appropriate. Between study heterogeneity was examined using the I^2^ statistic and meta-regression.

**Results:** Of 3,316 screened records, 63 were included. In randomly selected samples from the general population, the summary prevalence estimate was 1.3% (95% confidence intervals, CI 1.0 to 1.8%, I^2^ 41.5%, 3 studies) in countries with higher levels of development and 3.9% (95% CI 2.2 to 6.7, I^2^ 89.2%, 3 studies) in countries with lower levels. Prevalence estimates were similar in women and men (p=0.47). In clinic-based samples prevalence estimates were higher, except in asymptomatic patients (0.8%, 95% CI 0.4 to 1.4, I^2^ 0.0%, 3 studies). Summary prevalence estimates were: pregnant women 0.9% (95% CI 0.6 to 1.4%, I^2^ 0%, 4 studies); MSM in the community 3.2% (95% CI 2.1 to 5.1, I^2^ 78.3%, 5 studies); FSW in the community 15.9% (95% CI 13.5 to 18.9, I^2^ =79.9%, 4 studies).

**Discussion:** This systematic review can inform testing guidelines for *M. genitalium* infection. The low estimated prevalence of *M. genitalium* in the general population, pregnant women and asymptomatic attenders at clinics does not support expansion of testing to asymptomatic people in these groups.

Registration Numbers

PROSPERO: CRD42015020420

## Introduction

*Mycoplasma genitalium* is a cause of non-gonococcal non-chlamydial urethritis in men and cervicitis in women,^1-3^ and is reported to be associated with pelvic inflammatory disease, infertility and preterm birth.^4^ *M. genitalium* was first isolated in the early 1980s in men with non-gonococcal urethritis (NGU)^5^ but, owing to difficulties in detecting the microorganism by culture, most research on *M. genitalium* has been done since the development of nucleic acid amplification tests (NAAT) in the early 1990s.^1^ In populations studied in healthcare settings, *M. genitalium* has been detected in substantial proportions of study participants.^1^ ^2^ Based on these studies, routine testing has been suggested to detect and treat *M. genitalium* in asymptomatic attenders in healthcare settings^6^ and the recommendation has also been extended to low-risk general populations.^7^ Multiplex NAAT are being used increasingly to detect multiple sexually transmitted pathogens,^8^ ^9^ increasing pressure for their routine use.

Criteria for assessing the appropriateness of screening for a disease in the population include requirements that the disease is an important public health problem and that screening has been shown to do more good than harm.^10^ The prevalence of *M. genitalium* in asymptomatic populations has not been ascertained systematically. Non-systematic reviews have reported prevalence estimates ranging from zero to 0.7% to 3.3% in the general population^1^ and from zero to 20% in a range of female study populations described as ‘low-risk’.^11^ The objectives of this study were to systematically review the literature about the prevalence of *M. genitalium* in the general population and in specific groups (men who have sex with men, MSM, female sex workers, FSW, pregnant women, consecutively enrolled attenders in clinics).

## Methods

We followed a predefined review protocol.^12^ This report presents the findings of the first of three review questions (prevalence of *M. genitalium*). Two other review questions (incidence and persistence of untreated *M. genitalium* infection) will be addressed in a separate report. We report the findings using the Preferred Reporting Items for Systematic Reviews and Meta-Analyses (PRISMA, checklist in online Appendix, text S1).^13^

### Eligibility Criteria

We included studies that provided an estimate of the prevalence of *M. genitalium* infection in urogenital or rectal samples from women and men older than 13 years in any country from 1991 onwards, when the first NAAT was described.^1^ We included studies conducted amongst people in the general population or amongst attenders at healthcare settings. Eligible study designs were cross-sectional studies, cohort studies and baseline data in randomised controlled trials, published as full papers, abstracts or conference posters. We excluded laboratory studies, studies restricted to people with a specific condition, e.g., men with urethritis, women with abnormal cervical smears and women with pregnancy complications. To reduce small study effects, we restricted the review to larger studies that tend to use more rigorous study methodology and are at lower risk of bias.^14^ After screening titles and abstracts, we determined that inclusion of studies with 500 participants or more would result in at least 20 studies in the review.

### Information sources and search strategy

We searched Medline, Embase, African Index Medicus, IndMED, and LILACS databases from 1^st^ January 1991 to 12^th^ July 2016 without language restrictions. The full search strategy for Medline and Embase is provided in the online Appendix (Text S2). The other databases were searched using only the term “*Mycoplasma genitalium*.” We used Endnote (version 7, Thomson Reuters) to import, de-duplicate and manage retrieved records.

### Study selection

Two reviewers independently screened the identified records using pre-piloted checklists to assess eligibility, first of abstracts and titles and then of full text records. Differences were resolved by discussion or adjudication by a third reviewer. When multiple records reported on the same study population, we defined a primary record to represent the study, based on a combination of the following factors: description as a main paper by the authors, most detailed report of methods, prevalence reported as the main result and date of publication.

### Data collection process and data items

Two researchers extracted data independently for every included study, using a piloted extraction form in an online database (Research Electronic Data Capture, REDCap, Vanderbilt University, Tennessee). We resolved differences by discussion. The data extraction form included items about: study design, demographic characteristics, sample size, methods of participant selection and specimen collection, response rates, number of infected participants and number tested and reported prevalence estimates (with 95% confidence intervals, CI) overall and for prespecified subgroups.

We also recorded a measure of the level of development of the country in which the study was done using the human development index (HDI) 2015 HDI-dataset,^15^ which we categorised as higher (combining very high and high) or lower (medium and low). We defined studies *a priori* as ‘general population’ if they used any method to draw a random sample from the population of a whole country or a region, or as ‘community based’ if participants were enrolled outside healthcare settings but used non-random methods such as convenience sampling, snowball, or respondent driven sampling. Studies conducted in healthcare settings were coded according to their study population: clinic attenders, pregnant women, MSM and FSW. Studies that had enrolled participants from both healthcare settings and the community and did not stratify results were coded as clinic-based studies. We labelled studies according to the country in which the fieldwork was done and assigned numbers after the country name if there was more than one study from the same country. We generated separate strata within studies if they included participants from more than one country or from more than one relevant population subgroup, e.g. MSM and heterosexual adults.

### Risk of bias in individual studies

To evaluate the individual studies, we adapted an instrument from another systematic review of studies of *Chlamydia trachomatis* prevalence (online Appendix, text S3).^16^ Two reviewers independently assessed each item as being at high, low or uncertain risk of bias. Differences were resolved by discussion.

### Summary measure and synthesis of results

The outcome was the estimated prevalence (and 95% CI), defined as the number of specimens with a positive *M. genitalium* test result divided by the number of eligible participants with a valid test result. Where possible, we confirmed the published values using raw numbers reported in the publication. In studies that reported weighted prevalence estimates and confidence intervals or where raw numbers were not available, we used the information reported by the authors. We calculated survey response rates, whenever possible, by dividing the number of participants tested by the number of eligible people asked to participate.

We initially examined the estimates of *M. genitalium* prevalence visually in forest plots. We stratified studies, based on a previous study showing factors that contribute to heterogeneity in estimates of *C. trachomatis* prevalence,^16^ by: sampling method (random sample of the general population, community-based, or clinic-based); study population (general population, pregnant women, MSM, FSW); HDI (higher or lower); and, where reported, by sex and age of participants as under 25 years or 25 years and older.

We used the I^2^ statistic to assess heterogeneity that was not due to random variation.^17^ Heterogeneity was considered moderate or high when I^2^ was greater than 50% or 75% respectively. We used random effects meta-analysis to combine prevalence estimates where appropriate, assuming that, even when results were stratified, there might be real differences in *M. genitalium* prevalence between studies. We log-transformed the prevalence estimates and 95% CI before meta-analysis and back-transformed the summary average prevalence (and 95% CI) to the natural scale. We did not conduct meta-analysis on the logit scale because the log odds and confidence intervals could not be obtained from studies that reported weighted prevalence estimates. We calculated a prediction interval to provide information about the likely range of prevalence in future studies done in similar study populations if more than three studies contributed to a subgroup and if between-study heterogeneity was low (I^2^<50%).^18^ We did a meta-regression analysis to examine possible factors (HDI, use of probability sampling, sample size, response rate, sex and use of adequate sample and target populations) contributing to heterogeneity in general population and clinic-based studies. Analyses were done using the ‘metan’ and ‘metareg’ commands in Stata (Stata 13, Stata Corp, Austin, Texas, USA).

## Results

### Search results

We screened the titles and abstracts of 3,316 unique records published after 1991 and the full text of 833 studies (Figure S1 in online Appendix). A total of 63 records was included with participants who were; sampled at random from the general population^19-24^ or using alternative community-based methods,^25-29^ MSM and male-to-female transgendered,^30-35^ FSW,^36-40^ and pregnant women.^41-44^ Of these, 37 studies included patients attending healthcare settings.^8^ ^45-80^ We report results using the country name and number of the study or subgroup within a study.

Table 1 shows that most characteristics of included studies were similar to those of studies excluded because the sample size was below 500 (details in Table S1, online Appendix). The distribution of included and excluded studies was broadly similar. Eight of the excluded studies included participants from the community, but all studies that used probability-based sampling methods were included.

**Table 1.**
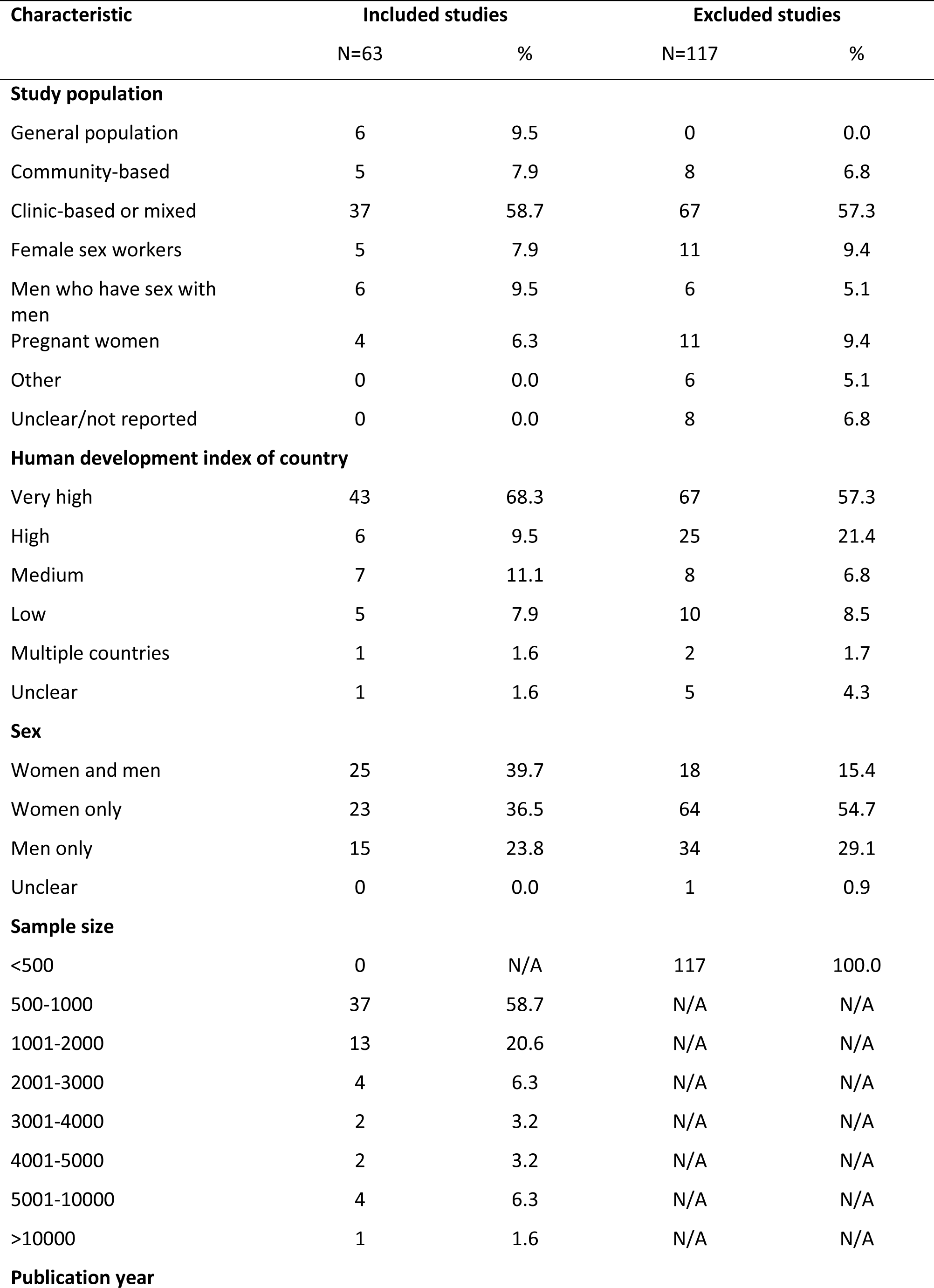
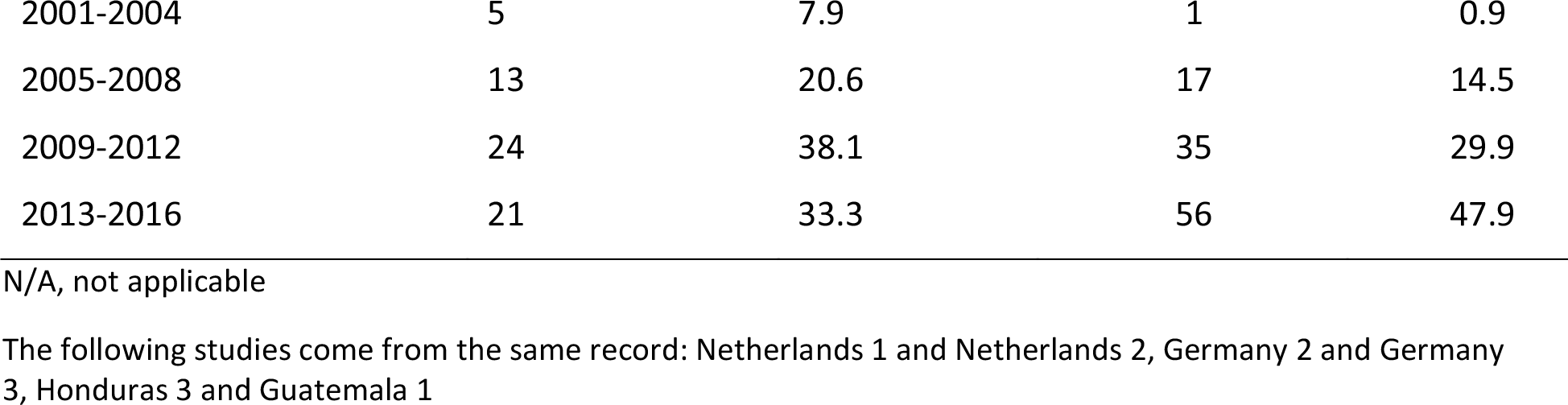
Characteristics of included and excluded studies

| Characteristic | Included studies |  | Excluded studies |  |
| --- | --- | --- | --- | --- |
|  | N=63 | % | N=117 | % |
| **Study population** |  |  |  |  |
| General population | 6 | 9.5 | 0 | 0.0 |
| Community-based | 5 | 7.9 | 8 | 6.8 |
| Clinic-based or mixed | 37 | 58.7 | 67 | 57.3 |
| Female sex workers | 5 | 7.9 | 11 | 9.4 |
| Men who have sex with men | 6 | 9.5 | 6 | 5.1 |
| Pregnant women | 4 | 6.3 | 11 | 9.4 |
| Other | 0 | 0.0 | 6 | 5.1 |
| Unclear/not reported | 0 | 0.0 | 8 | 6.8 |
| **Human development index of country** |  |  |  |  |
| Very high | 43 | 68.3 | 67 | 57.3 |
| High | 6 | 9.5 | 25 | 21.4 |
| Medium | 7 | 11.1 | 8 | 6.8 |
| Low | 5 | 7.9 | 10 | 8.5 |
| Multiple countries | 1 | 1.6 | 2 | 1.7 |
| Unclear | 1 | 1.6 | 5 | 4.3 |
| **Sex** |  |  |  |  |
| Women and men | 25 | 39.7 | 18 | 15.4 |
| Women only | 23 | 36.5 | 64 | 54.7 |
| Men only | 15 | 23.8 | 34 | 29.1 |
| Unclear | 0 | 0.0 | 1 | 0.9 |
| **Sample size** |  |  |  |  |
| <500 | 0 | N/A | 117 | 100.0 |
| 500-1000 | 37 | 58.7 | N/A | N/A |
| 1001-2000 | 13 | 20.6 | N/A | N/A |
| 2001-3000 | 4 | 6.3 | N/A | N/A |
| 3001-4000 | 2 | 3.2 | N/A | N/A |
| 4001-5000 | 2 | 3.2 | N/A | N/A |
| 5001-10000 | 4 | 6.3 | N/A | N/A |
| >10000 | 1 | 1.6 | N/A | N/A |
| **Publication year** |  |  |  |  |
| before 2000 | 0 | 0.0 | 8 | 6.8 |
| 2001-2004 | 5 | 7.9 | 1 | 0.9 |
| 2005-2008 | 13 | 20.6 | 17 | 14.5 |
| 2009-2012 | 24 | 38.1 | 35 | 29.9 |
| 2013-2016 | 21 | 33.3 | 56 | 47.9 |

N/A, not applicable
The following studies come from the same record: Netherlands 1 and Netherlands 2, Germany 2 and Germany 3, Honduras 3 and Guatemala 1

### Risk of bias in individual studies

No study was at low risk of bias in all domains (Figure S2). The studies at lowest risk of bias were those that used probability sampling in the general population. Only one study compared responders and non-responders and that study found differences between these groups.^24^ Reporting of complete results, including CIs and baseline data, was considered adequate in 22 studies.

### Studies in the general population and community

We included 11 studies, six of which were in countries with higher HDI (Denmark 1,^23^ Great Britain 2 and Great Britain 4,^24^ ^25^ Norway 4,^26^ Russian Federation 3^27^ and USA 2,^19^ N=13,331) and five in countries with a lower (Honduras 1,^20^ Vietnam 1,^21^ Kenya 1,^28^ Madagascar 1,^29^ Tanzania 1,^22^ N=4,978) HDI (Figure 1, Table S2 in online Appendix).

**Figure 1.**
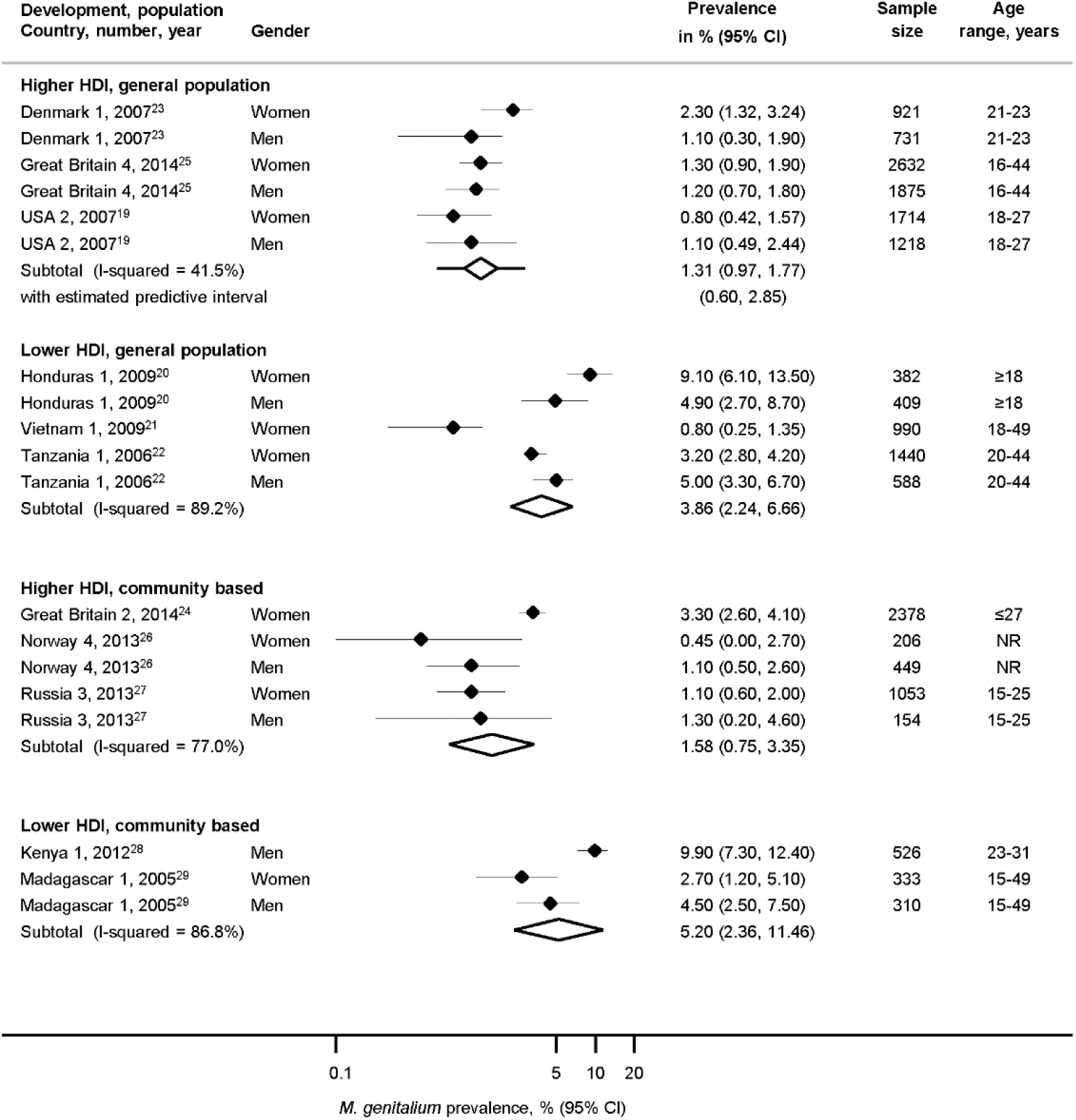
Estimated prevalence of *Mycoplasma genitalium* in randomly selected samples of the general population or in other community-based samples, by human development index. CI, confidence interval; HDI, human development index. Solid diamond and lines show the point estimate and 95% confidence intervals for each study. The diamond shows the point estimate and 95% confidence intervals of the summary estimate. Lines extending from the diamond show the prediction interval. The prevalence estimates are plotted on a logarithmic scale.

The summary average general population prevalence of *M. genitalium* in three studies in countries with higher HDI was 1.3% (95% CI 1.0 to 1.8%, prediction interval 0.6, 2.9%, I^2^ 41.5%, N=9,091, Figure 1), with low between-study heterogeneity in three studies (one region in Denmark 1,^23^ or the whole population in Great Britain 4 and USA 2).^19^ ^24^ In three studies in higher HDI countries that enrolled participants using convenience sampling methods from sub-national communities (N=4,240, Table S2), between-study heterogeneity was higher than in the studies that used random sampling methods but the summary average prevalence was similar (1.6%, 95% 0.8 to 3.4%, I^2^ 77.0%).^25-27^ Amongst adults under 25 years, average *M. genitalium* prevalence was 1.7% (95% CI 1.0 to 2.6%, I^2^ 80.3%) in women and 0.3% in men (0.1 to 1.4%, I^2^ 91.3%) (Figure S3 in online Appendix). There were too few estimates from adults aged 25 years and over to compare estimates between women and men.

The surveys from five countries with lower HDI enrolled very different populations *M. genitalium* prevalence estimates were more variable (Figure 1, Table S2 in online Appendix).^20-22^ ^28^ ^29^ The summary estimate of prevalence in three studies that used probability sampling was 3.9% (2.2 to 6.7, I^2^ 89.2%) and, in two studies that used other methods to enrol participants from community settings, 5.2% (2.4 to 11.5, I^2^ 86.8%).

In a meta-regression analysis that compared characteristics of all studies in adults in the general population, there was some statistical evidence to suggest higher estimates of *M. genitalium* prevalence in countries with lower than higher HDI (difference 3.1%, 95% CI −0.1 to 6.3%, p=0.057) but no statistical evidence of a difference by sex (0.9%, 95% CI −1.6 to 3.3% p=0.47) or for other study related variables that were examined (Table S3 in online appendix).

### Pregnant women in antenatal clinics and women in the general population

We included four studies in pregnant women before 14 weeks gestation, all in countries with higher HDI (N=3,472, age range 16 to 48 years; France 2,^44^ Great Britain 1,^41^ Japan 1,^42^ and USA 5,^43^ Figure 2, Table S4 in online Appendix). The combined average prevalence was 0.9% (95% CI 0.6 to 1.4%, prediction interval 0.4, 2.3%, I^2^ 0%). The estimated prevalence was slightly lower than in the three studies in women in the general population (1.4%, 95% CI 0.8 to 2.4%, I^2^ 73.4%) but confidence intervals overlapped.

**Figure 2.**
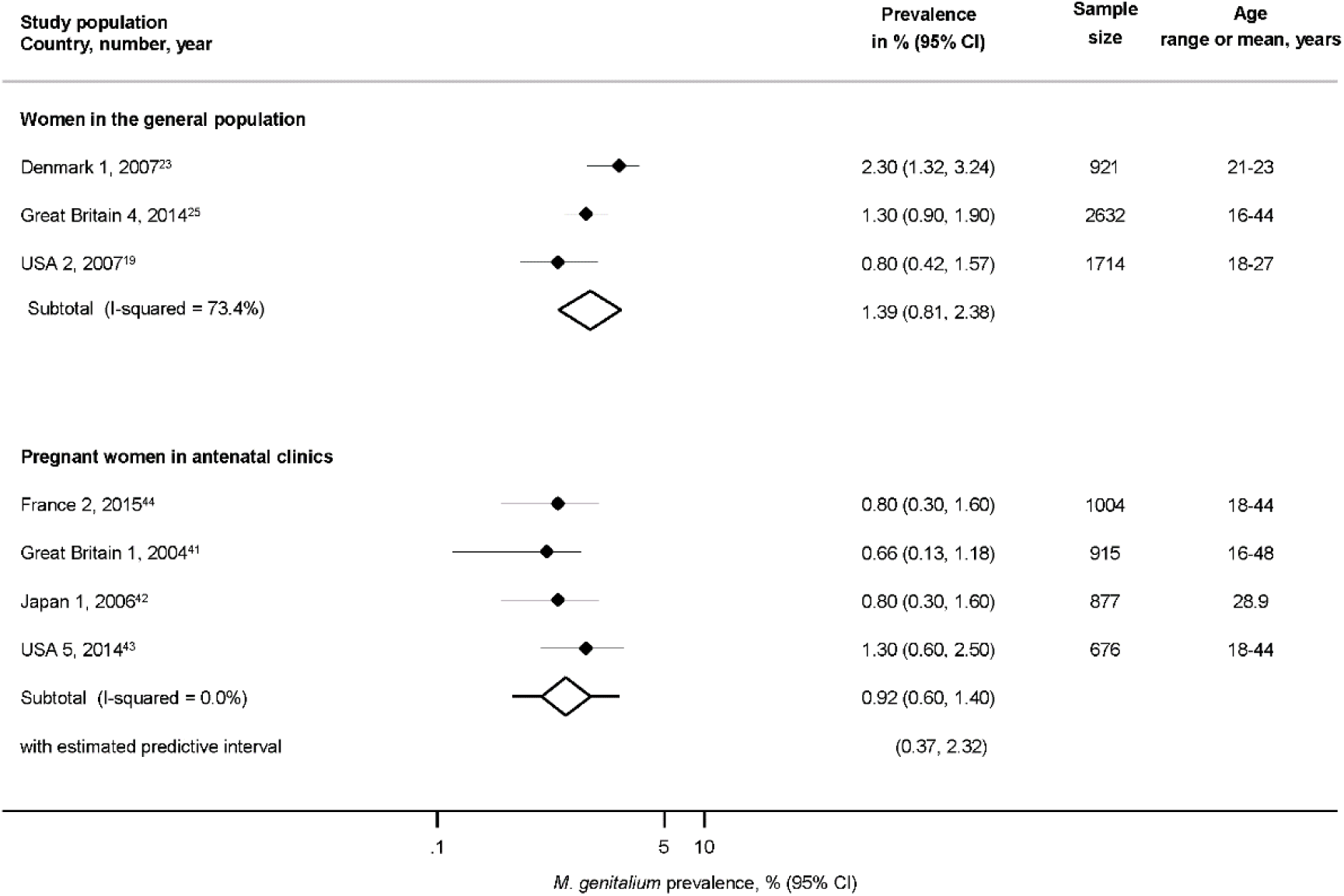
Estimated prevalence of *Mycoplasma genitalium* in pregnant women in antenatal clinics and in randomly selected samples of women in the general population. CI, confidence interval. Solid diamond and lines show the point estimate and 95% confidence intervals for each study. The diamond shows the point estimate and 95% confidence intervals of the summary estimate. Lines extending from the diamond show the prediction interval. The prevalence estimates are plotted on a logarithmic scale.

### MSM and FSW in community-based and clinic-based studies

Five studies from four records enrolled MSM from the community (Figure 3, Table S4 in online Appendix) in specific areas in; Australia 2,^30^ El Salvador 1,^31^ Guatemala 1 and Honduras 3,^32^ and Nicaragua 1,^33^ (N=3,012). The summary average prevalence in these studies was 3.2% (95% CI 2.1 to 5.1%, I^2^ 78.3%) with moderate between-study heterogeneity. The summary average estimate of *M. genitalium* prevalence in MSM enrolled from clinics in Germany 3,^55^ the Netherlands 2,^54^ Norway 5^35^ and USA 3^34^ was 3.7% (95% CI 2.4 to 5.6%, I^2^ 78.5%).

**Figure 3.**
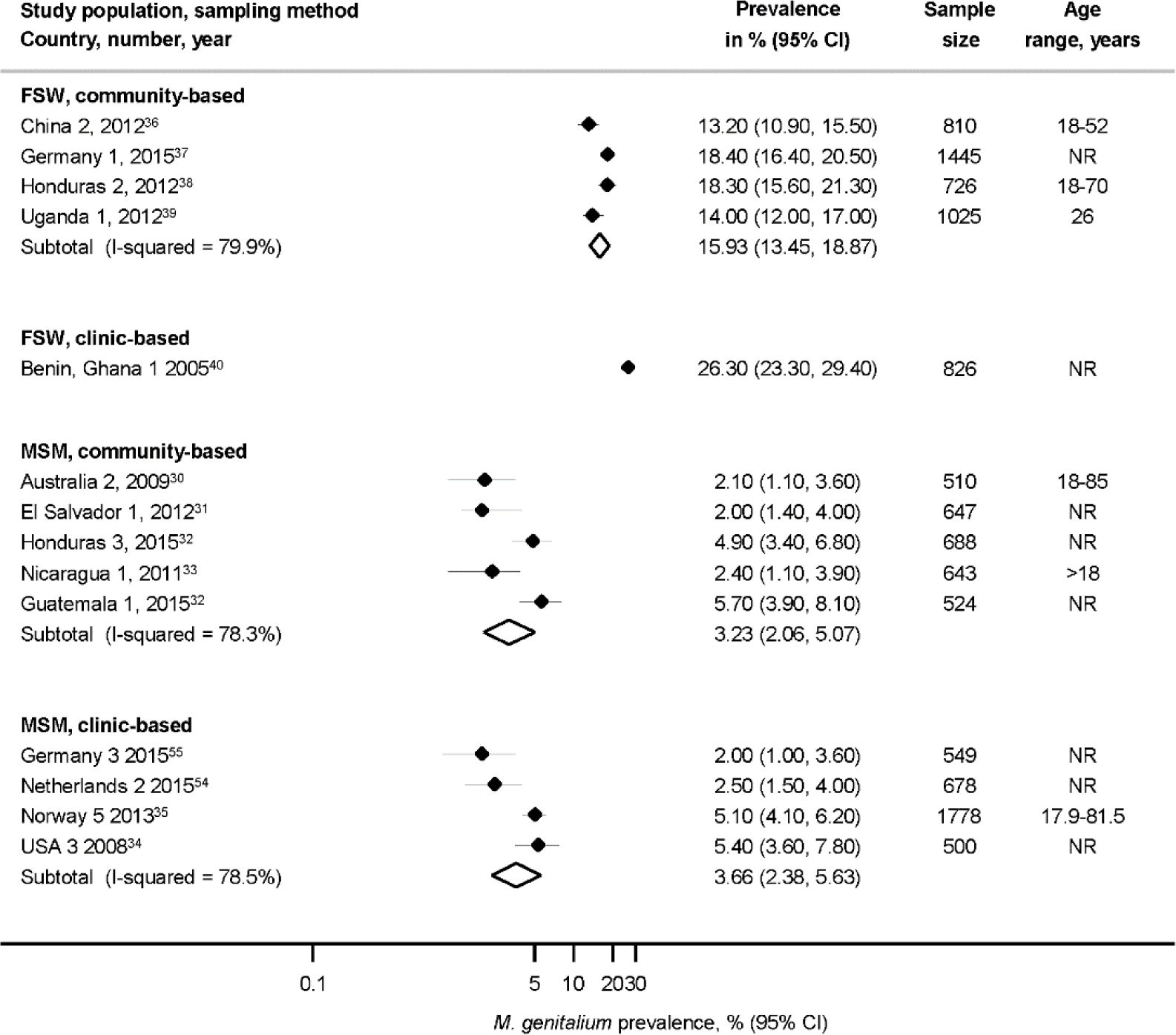
Estimated prevalence of *Mycoplasma genitalium* in community-based and clinic-based samples of men who have sex with men and female sex workers CI, confidence interval; FSW, female sex workers; MSM, men who have sex with men; NR, not reported. Solid diamond and lines show the point estimate and 95% confidence intervals for each study. The diamond shows the point estimate and 95% confidence intervals of the summary estimate. Lines extending from the diamond show the prediction interval. The prevalence estimates are plotted on a logarithmic scale.

Studies that enrolled FSW were more variable. Estimated *M. genitalium* prevalence in the community-based studies in specific areas in southwest China 2,^36^ northern Germany 1,^37^ Honduras 2,^38^ and Uganda 1^39^ was 15.9% (95% CI 13.5 to 18.9%, N=4,006, I^2^ 79.9%). In one study that enrolled FSW from a clinic in Benin and Ghana 1,^40^ the estimated prevalence was higher (26.3%, 95% CI 23.3 to 29.4%).^40^

### Clinic-based studies, unselected populations

We included 37 studies from 14 countries, of which 24 were from Australia, Great Britain, Norway, South Korea and Sweden (Table S5, Figure S4 in online Appendix). Estimates of *M. genitalium* were very heterogeneous (I^2^ >95%), except for studies that only included patients without symptoms,^8^ ^50^ ^64^ (N=2889, 0.8%, 95% CI 0.4 to 1.4%, I^2^ 0%). Most study populations included patients both with and without symptoms. The lowest estimate was reported in a fertility clinic population in Venezuela 1^78^ (N=3,358, 0.6%, 95% CI 0.3 to 0.9%). The highest estimates were from patients attending youth clinics in Russia 1(N=9,208, 12.6%, 95% CI 11.9 to 13.3%), women attending primary health care clinics in Limpopo province, South Africa 1^63^ (N=601, 8.7%, 95% CI 6.4 to 10.9%) and STI clinic attenders in Sweden 8^73^ (N=679, 7.4% 95% CI 5.5 to 9.6%) and the USA 4^77^ (N=1,090, 7.7%, 95% CI 6.2 to 9.4%).

## Discussion

### Main Findings

In large nationally representative surveys conducted in very highly developed countries the summary average prevalence estimate of *M. genitalium* was 1.3% (95% CI 1.0 to 1.8%,3 studies, I^2^ 41.5%) in adults aged 16 to 44 years with no statistical evidence of a difference between men and women (p=0.47). Summary prevalence estimates in specific subpopulations were: pregnant women 0.9% (0.6 to 1.4%), MSM in community samples 3.2% (2.1 to 5.1%, 5 studies, I^2^ 78.3%) and MSM in clinic-based samples 3.7% (2.4 to 5.6%, 4 studies, I^2^ 78.5%). Prevalence estimates were higher in FSW, ranging from 13.2% in one community-based study to 26.3% in one clinic-based study. In clinic-based surveys, prevalence estimates varied from 0.8% (95% CI 0.4 to 1.4%, 3 studies, I^2^ 0%) in patients with no symptoms to 12.6% (95% CI 11.9 to 13.3%) in one study in Russia.

### Strengths and limitations

The broad search strategy is a strength of this review. It allowed for identification of a wide range of different studies and it is unlikely that we missed large studies. The *a priori* defined inclusion criteria allowed a clear selection process for the detected studies and duplicate screening and data extraction prevented data entry errors. By including only studies with 500 participants or more, we aimed to reduce the influence of small study biases that can distort results. This strategy included all studies that used methods to select random samples of the general population and provided summary estimates with little heterogeneity for general population samples in very highly developed countries, pregnant women and asymptomatic people attending outpatient healthcare settings. Although we explored between-study heterogeneity using meta-regression analysis, we did not identify factors that could explain a substantial proportion of the heterogeneity. A further limitation to interpretation is that we could not provide prediction intervals for most subgroups because the method used to estimate prediction intervals is not suited to situations with few studies or where heterogeneity or the risk of bias are high.^18^ Finally, we could not assess an earlier finding, in surveys of chlamydia prevalence,^16^ that lower response rates are associated with higher prevalence estimate because very few studies reported these results. Amongst studies that reported response rates we did not find an association with *M. genitalium* prevalence (Table S3 in online Appendix).

### Interpretation and comparison with other studies

To our knowledge, this is the first systematic review assessing the prevalence of *M. genitalium* in different population groups including those outside healthcare settings. Our findings suggest that *M. genitalium* might be less prevalent than *C. trachomatis* in the general population but comparison is not straightforward. In a systematic review of population-based surveys of *C. trachomatis*, estimated prevalence in adults <27 years in high-income countries was 4.3% (95% CI 3.6 to 5.0%, I^2^ 0%) in women and 3.6% (95% CI 2.8 to 4.4%, I^2^ 6.2%) in men,^16^ compared with our summary estimates of less than 2% for *M. genitalium* in women and men <25 years old. Within studies that tested for both pathogens, prevalence estimates for *M. genitalium* and *C. trachomatis* were similar in Great Britain,^24^ but higher for *C. trachomatis* than *M. genitalium* in Denmark^23^ and the USA.^19^ It is, however, possible that *M. genitalium* prevalence has been underestimated because the sensitivity of NAATs is lower than previously believed.^9^ In general, age differences seem less marked amongst women for *M. genitalium* than for *C. trachomatis*, where prevalence after age 25 years is much lower than in younger women. Age-specific patterns of *M. genitalium* were, however, difficult to discern with certainty, largely because population-based studies that provided age-stratified estimates used non-comparable age groups and only two had estimates for participants older than 25 years (Norway 4 and Great Britain 4 studies).

In clinic-based surveys, participant selection methods and characteristics differed substantially between different types of clinics and countries. *M. genitalium* prevalence estimates were consistent and comparable (or even lower) than in general population-based surveys in studies that only enrolled asymptomatic patients or pregnant women in antenatal clinics. Amongst MSM, estimated *M. genitalium* prevalence was similar in community-based and clinic-based studies.

### Implications for clinical practice, policy and research

This systematic review provides evidence about the prevalence of *M. genitalium* that can be used in mathematical modelling studies and to inform testing guidelines for infection. The trend for molecular diagnostic tests to include targets that identify multiple sexually transmitted pathogens means that testing for asymptomatic *M. genitalium* infection will become more widespread. In clinical settings, systematic reviews of the prevalence of *M. genitalium* in symptomatic non-gonococcal non-chlamydial urethritis and of resistance to macrolides and fluoroquinolones would further help to refine clinical treatment guidelines for *M. genitalium* infections. The absence of randomised controlled trials that demonstrate a clinical benefit of screening and the increasing prevalence of resistance to azithromycin are reasons for restricting widespread testing for *M. genitalium*.^81^ The low estimated prevalence of *M. genitalium* in the general population, in pregnant women and in asymptomatic attenders in health care settings and absence of a clearly defined age group at higher risk of infection do not provide strong support for the appropriateness of universal or age-based screening programmes for *M. genitalium* in these population groups.

### Key messages

- Routine screening for *Mycoplasma genitalium* infection has been proposed, but prevalence rates are not well established
- In samples from the general population, the summary prevalence estimate is 1.3% in countries with higher development and 3.9% in countries with lower development
- *M. genitalium* prevalence in the general population and differences in prevalence by age appear to be less than for *Chlamydia trachomatis*
- The low prevalence estimates in the general population, pregnant women and asymptomatic clinic-based patients do not support universal screening for *M. genitalium*

## Author contributions

Conceived and designed the review; NL, LB, MC, PS, DE. Screened titles, abstracts and full text; LB, MC, DE, HA, GL, NL. Extracted data; LB, MC, DE, HA, G-RL. Analysed data; LB, MC, FH, MG. Wrote the first draft; LB, MC. Revised the paper; NL, MG, FH, DE, PS, HA. Approved the final version; LB, MC, DE, MG, FH, PS, HA, NL.

## Author contributions

We would like to thank Georgia Salanti for her advice about meta-analysis.

## References

1. Taylor-Robinson D, Jensen JS. Mycoplasma genitalium: from Chrysalis to multicolored butterfly. Clin Microbiol Rev 2011;24(3):498–514.

2. Manhart LE, Kay N. Mycoplasma genitalium: Is it a sexually transmitted pathogen? Curr Infect Dis Rep 2010;12(4):306–13.

3. Hjorth SV, Bjornelius E, Lidbrink P, et al. Sequence-based typing of Mycoplasma genitalium reveals sexual transmission. J Clin Microbiol 2006;44(6):2078–83.

4. Lis R, Rowhani-Rahbar A, Manhart LE. Mycoplasma genitalium Infection and Female Reproductive Tract Disease: A Meta-analysis. Clin Infect Dis 2015;61(3):418–26.

5. Tully JG, Taylor-Robinson D, Cole RM, et al. A newly discovered mycoplasma in the human urogenital tract. Lancet 1981;1(8233):1288–91.

6. Manhart LE. Has the time come to systematically test for Mycoplasma genitalium? Sex Transm Dis 2009;36(10):607–8.

7. McGowin CL, Rohde RE, Redwine G. Epidemiological and clinical rationale for screening and diagnosis of Mycoplasma genitalium infections. Clin Lab Sci 2014;27(1):47–52.

8. Kim SJ, Lee DS, Lee SJ. The prevalence and clinical significance of urethritis and cervicitis in asymptomatic people by use of multiplex polymerase chain reaction. Korean Journal of Urology 2011;52(10):703–08.

9. Gaydos CA. Mycoplasma genitalium: Accurate Diagnosis Is Necessary for Adequate Treatment. J Infect Dis 2017; 216(Suppl_2):S406–S11.

10. Raffle A, Gray M. Screening: Evidence and practice. Oxford: Oxford University Press 2007.

11. McGowin CL, Anderson-Smits C. Mycoplasma genitalium: an emerging cause of sexually transmitted disease in women. PLoS Pathog 2011;7(5):e1001324.

12. Low N, Cina M, Baumann L, et al., Mycoplasma genitalium infection: prevalence, incidence and persistence. PROSPERO 2015:CRD42015020420 Systematic review protocol. (accessed 09.05.2017.).

13. Moher D, Liberati A, Tetzlaff J, et al. Preferred Reporting Items for Systematic Reviews and Meta-Analyses: The PRISMA Statement. PLoS Med 2009;6(7):e1000097.

14. Vandenbroucke JP, Von EE, Altman DG, et al. Strengthening the Reporting of Observational Studies in Epidemiology (STROBE): explanation and elaboration. PLoS Med 2007;4(10):e297.

15. United Nations Development Programme. Human development report 2016: Human development for everyone. New York, USA, 2016:198–201.

16. Redmond SM, Alexander-Kisslig K, Woodhall SC, et al. Genital chlamydia prevalence in Europe and non-European high income countries: systematic review and meta-analysis. PLoS One 2015;10(1):e0115753.

17. Higgins JP, Thompson SG. Quantifying heterogeneity in a meta-analysis. Stat Med 2002;21(11):1539–58.

18. Riley RD, Higgins JP, Deeks JJ. Interpretation of random effects meta-analyses. BMJ 2011;342:d549.

19. Manhart LE, Holmes KK, Hughes JP, et al. Mycoplasma genitalium among young adults in the United States: an emerging sexually transmitted infection. Am J Public Health 2007;97(6):1118–25.

20. Paz-Bailey G, Morales-Miranda S, Jacobson JO, et al. High rates of STD and sexual risk behaviors among Garifunas in Honduras. J Acquir Immune Defic Syndr 2009;51 Suppl 1:S26–34.

21. Olsen B, Lan PT, Stalsby Lundborg C, et al. Population-based assessment of Mycoplasma genitalium in Vietnam--low prevalence among married women of reproductive age in a rural area. J Eur Acad Dermatol Venereol 2009;23(5):533–7.

22. Kapiga SH, Sam NE, Mlay J, et al. The epidemiology of HIV-1 infection in northern Tanzania: results from a community-based study. AIDS Care 2006;18(4):379–87.

23. Andersen B, Sokolowski I, Ostergaard L, et al. Mycoplasma genitalium: prevalence and behavioural risk factors in the general population. Sex Transm Infect 2007;83(3):237–41.

24. Sonnenberg P, Ison CA, Clifton S, et al. Epidemiology of Mycoplasma genitalium in British men and women aged 16-44 years: Evidence from the third National Survey of Sexual Attitudes and Lifestyles (Natsal-3). Int J Epidemiol 2015;44(6):1982–94.

25. Oakeshott P, Aghaizu A, Hay P, et al. Is Mycoplasma genitalium in women the “New Chlamydia?” A community-based prospective cohort study. Clin Infect Dis 2010;51(10):1160–6.

26. Jensen AJ, Kleveland CR, Moghaddam A, et al. Chlamydia trachomatis, Mycoplasma genitalium and Ureaplasma urealyticum among students in northern Norway. J Eur Acad Dermatol Venereol 2013;27(1):e91–e96.

27. Shipitsyna E, Khusnutdinova T, Ryzhkova O, et al. Prevalence of Sexually Transmitted Infections in Young People in St. Petersburg, Russia, as Determined Using Self-Collected Non-Invasive Specimens. Sex Transm Infect 2013;89:A94–A94.

28. Mehta SD, Gaydos C, Maclean I, et al. The effect of medical male circumcision on urogenital Mycoplasma genitalium among men in Kisumu, Kenya. Sex Transm Dis 2012;39(4):276–80.

29. Leutscher P, Jensen JS, Hoffmann S, et al. Sexually transmitted infections in rural Madagascar at an early stage of the HIV epidemic: a 6-month community-based follow-up study. Sex Transm Dis 2005;32(3):150–5.

30. Bradshaw CS, Fairley CK, Lister NA, et al. Mycoplasma genitalium in men who have sex with men at male-only saunas. Sex Transm Infect 2009;85(6):432–5.

31. Creswell J, Guardado ME, Lee J, et al. HIV and STI control in El Salvador: results from an integrated behavioural survey among men who have sex with men. Sex Transm Infect 2012;88(8):633–8.

32. Ham D, Northbrook SY, Morales-Miranda S, et al. HIV and STIs among transgendered populations: Four country survey from central America [abstract]. Top Antivir Med 2015;23(E-1):475–76.

33. Hernandez F, Arambu N, Alvarez B, et al. High incidence of HIV and low HIV prevention coverage among men who have sex with men in Managua, Nicaragua. Sex Transm Infect 2011;87:A146.

34. Francis SC, Kent CK, Klausner JD, et al. Prevalence of rectal Trichomonas vaginalis and Mycoplasma genitalium in male patients at the San Francisco STD clinic, 2005-2006. Sex Transm Dis 2008;35(9):797–800.

35. Reinton N, Moi H, Olsen AO, et al. Anatomic distribution of Neisseria gonorrhoeae, Chlamydia trachomatis and Mycoplasma genitalium infections in men who have sex with men. Sex Health 2013;10(3):199–203.

36. Xiang Z, Yin YP, Shi MQ, et al. Risk factors for Mycoplasma genitalium infection among female sex workers: a cross-sectional study in two cities in southwest China. BMC Public Health 2012;12:414.

37. Jansen K, Bremer V, Steffen G, et al. High prevalence of genital infections with mycoplasma genitalium in female sex workers reached at their working place in Germany: The STI-outreach-study. Sex Transm Infect 2015;91:A149–A50.

38. Johnston LG, Paz-Bailey G, Morales-Miranda S, et al. High prevalence of Mycoplasma genitalium among female sex workers in Honduras: implications for the spread of HIV and other sexually transmitted infections. Int J STD AIDS 2012;23(1):5–11.

39. Vandepitte J, Muller E, Bukenya J, et al. Prevalence and correlates of Mycoplasma genitalium infection among female sex workers in Kampala, Uganda. J Infect Dis 2012;205(2):289–96.

40. Pepin J, Labbe AC, Khonde N, et al. Mycoplasma genitalium: an organism commonly associated with cervicitis among west African sex workers. Sex Transm Infect 2005;81(1):67–72.

41. Oakeshott P, Hay P, Taylor-Robinson D, et al. Prevalence of Mycoplasma genitalium in early pregnancy and relationship between its presence and pregnancy outcome. BJOG 2004;111(12):1464–7.

42. Kataoka S, Yamada T, Chou K, et al. Association between preterm birth and vaginal colonization by mycoplasmas in early pregnancy. J Clin Microbiol 2006;44(1):51–5.

43. Agger WA, Siddiqui D, Lovrich SD, et al. Epidemiologic factors and urogenital infections associated with preterm birth in a midwestern U.S. population. Obstet Gynecol 2014;124(5):969–77.

44. Peuchant O, Le Roy C, Desveaux C, et al. Screening for Chlamydia trachomatis, Neisseria gonorrhoeae, and Mycoplasma genitalium should it be integrated into routine pregnancy care in French young pregnant women? Diagn Microbiol Infect Dis 2015;82(1):14–9.

45. McKechnie ML, Hillman R, Couldwell D, et al. Simultaneous identification of 14 genital microorganisms in urine by use of a multiplex PCR-based reverse line blot assay. J Clin Microbiol 2009;47(6):1871–7.

46. Walker J, Fairley CK, Bradshaw CS, et al. 'The difference in determinants of Chlamydia trachomatis and Mycoplasma genitalium in a sample of young Australian women'. BMC Infect Dis 2011;11:35.

47. Lusk MJ, Konecny P, Naing ZW, et al. Mycoplasma genitalium is associated with cervicitis and HIV infection in an urban Australian STI clinic population. Sex Transm Infect 2011;87(2):107–9.

48. Bao T, Chen R, Zhang J, et al. Simultaneous detection of Ureaplasma parvum, Ureaplasma urealyticum, Mycoplasma genitalium and Mycoplasma hominis by fluorescence polarization. J Biotechnol 2010;150(1):41–3.

49. Sednaoui P, Nassar N, Allemelou G, et al. Evaluation of the Bio-rad Dx CT/NG/MG assay, a new real-time PCR test for the simultaneous detection of Chlamydia trachomatis, Neisseria gonorrhoeae and Mycoplasma genitalium. Clin Microbiol Infect 2011;17:S486.

50. Clarivet B, Picot E, Marchandin H, et al. Prevalence of Chlamydia trachomatis, Neisseria gonorrhoeae and Mycoplasma genitalium in asymptomatic patients under 30 years of age screened in a French sexually transmitted infections clinic. Eur J Dermatol 2014;24(5):611–16.

51. Jalal H, Delaney A, Bentley N, et al. Molecular epidemiology of selected sexually transmitted infections. Int J Mol Epidemiol Genet 2013;4(3):167–74.

52. Svenstrup HF, Dave SS, Carder C, et al. A cross-sectional study of Mycoplasma genitalium infection and correlates in women undergoing population-based screening or clinic-based testing for Chlamydia infection in London. BMJ Open 2014;4(2)

53. Slack R, Yavuz M, Elangasinghe M, et al. The prevalence of mycoplasma genitalium among male gum attendees using an in-house PCR on BD MAX platform. HIV Med 2014;15:11.

54. Van Der Veer C, Van Rooijen MS, De Vries HJC, et al. Trichomonas vaginalis and mycoplasma genitalium: Age-specific prevalence and disease burden in men attending a sexually transmitted infections clinic in Amsterdam, The Netherlands. Sex Transm Infect 2015;91:A148–A49.

55. Lallemand A, Bremer V, Jansen K, et al. Prevalence of mycoplasma genitalium in patients visiting HIV counselling institutions in north-rhine-westphalia, Germany (STI-hit study). Sex Transm Infect 2015;91:A150.

56. Moi H, Reinton N, Moghaddam A. Mycoplasma genitalium is associated with symptomatic and asymptomatic non-gonococcal urethritis in men. Sex Transm Infect 2009;85(1):15–8.

57. Moi H, Reinton N, Moghaddam A. Mycoplasma genitalium in women with lower genital tract inflammation. Sex Transm Infect 2009;85(1):10–4.

58. Nilsen E, Vik E, Roed MA. Low prevalence of Mycoplasma genitalium in patients examined for Chlamydia trachomatis. Tidsskr Nor Laegeforen 2011;131(22):2232–4.

59. Hartgill U, Kalidindi K, Molin SB, et al. Screening for Chlamydia trachomatis and Mycoplasma genitalium; is first void urine or genital swab best? Sex Transm Infect 2015;91(2):141.

60. Reinton N, Hjelmevoll SO, Haheim H, et al. Analysis of direct-to-consumer marketed Chlamydia trachomatis diagnostic tests in Norway. Sexual Health 2015;12(4):336–40.

61. Khryanin A, Reshetnikov O. The detection rate of chlamydia trachomatis and mycoplasma genitalium infections in std clinics in novosibirsk, Russian federation. Sex Transm Infect 2011;87:A101–A02.

62. Berle LM, Firsova N, Kalashnik A, et al. Chlamydia trachomatis, Mycoplasma genitalium and Ureaplasma urealyticum in clinical and non-clinical settings, Arkhangelsk Oblast, Russia. Int J STD AIDS 2012;23(11):781–84.

63. Hay B, Dubbink JH, Ouburg S, et al. Prevalence and macrolide resistance of mycoplasma genitalium in South African women. Sex Transm Dis 2015;42(3):140–42.

64. Choi JY, Cho IC, Lee GI, et al. Prevalence and associated factors for four sexually transmissible microorganisms in middle-aged men receiving general prostate health checkups: A polymerase chain reaction-based study in Korea. Korean Journal of Urology 2013;54(1):53–58.

65. Kim Y, Kim J, Lee KA. Prevalence of sexually transmitted infections among healthy Korean women: implications of multiplex PCR pathogen detection on antibiotic therapy. J Infect Chemother 2014;20(1):74–6.

66. Falk L, Fredlund H, Jensen JS. Tetracycline treatment does not eradicate Mycoplasma genitalium. Sex Transm Infect 2003;79(4):318–9.

67. Falk L, Fredlund H, Jensen JS. Symptomatic urethritis is more prevalent in men infected with Mycoplasma genitalium than with Chlamydia trachomatis. Sex Transm Infect 2004;80(4):289–93.

68. Jensen JS, Bjornelius E, Dohn B, et al. Comparison of first void urine and urogenital swab specimens for detection of Mycoplasma genitalium and Chlamydia trachomatis by polymerase chain reaction in patients attending a sexually transmitted disease clinic. Sex Transm Dis 2004;31(8):499–507.

69. Mellenius H, Boman J, Lundqvist EN, et al. [Mycoplasma genitalium should be suspected in unspecific urethritis and cervicitis. A study from Vasterbotten confirms the high prevalence of the bacteria]. Lakartidningen 2005;102(47):3538, 40–1.

70. Anagrius C, Lore B, Jensen JS. Mycoplasma genitalium: prevalence, clinical significance, and transmission. Sex Transm Infect 2005;81(6):458–62.

71. Jurstrand M, Jensen JS, Fredlund H, et al. Detection of Mycoplasma genitalium in urogenital specimens by real-time PCR and by conventional PCR assay. J Med Microbiol 2005;54(Pt 1):23–9.

72. Hogdahl M, Kihlstrom E. Leucocyte esterase testing of first-voided urine and urethral and cervical smears to identify Mycoplasma genitalium-infected men and women. Int J STD AIDS 2007;18(12):835–8.

73. Edberg A, Jurstrand M, Johansson E, et al. A comparative study of three different PCR assays for detection of Mycoplasma genitalium in urogenital specimens from men and women. J Med Microbiol 2008;57(Pt 3):304–9.

74. Bjartling C, Osser S, Persson K. Mycoplasma genitalium in cervicitis and pelvic inflammatory disease among women at a gynecologic outpatient service. Am J Obstet Gynecol 2012;206(6):476 e1–8.

75. Tobian AA, Gaydos C, Gray RH, et al. Male circumcision and Mycoplasma genitalium infection in female partners: a randomised trial in Rakai, Uganda. Sex Transm Infect 2014;90(2):150–4.

76. Manhart LE, Critchlow CW, Holmes KK, et al. Mucopurulent cervicitis and Mycoplasma genitalium. J Infect Dis 2003;187(4):650–7.

77. Hancock EB, Manhart LE, Nelson SJ, et al. Comprehensive assessment of sociodemographic and behavioral risk factors for Mycoplasma genitalium infection in women. Sex Transm Dis 2010;37(12):777–83.

78. Peralta-Arias RD, Chollett D, Del Gobbo A, et al. Identification of ureaplasma SPP, chlamydia trachomatis and mycoplasma genitalium using multiplex real time PCR in cervical swabs from women attending a fertility institute in venezuela. Fertil Steril 2013;100(3):S379.

79. Gesink D, Sarai Racey C, Seah C, et al. Mycoplasma genitalium in Toronto, Ont: Estimates of prevalence and macrolide resistance. Can Fam Physician 2016;62(2):e96–e101.

80. Leung A, Eastick K, Haddon LE, et al. Mycoplasma genitalium is associated with symptomatic urethritis. Int J STD AIDS 2006;17(5):285–8.

81. Golden MR, Workowski KA, Bolan G. Developing a Public Health Response to Mycoplasma genitalium. J Infect Dis 2017;216(suppl_2):S420–S26.

